# A metabolite sensor subunit of the Atg1/ULK complex regulates selective autophagy

**DOI:** 10.1101/2022.12.13.520293

**Authors:** A.S. Gross, R. Ghillebert, M. Schuetter, E. Reinartz, A. Rowland, M. Graef

## Abstract

Cells convert complex metabolic information into stress-adapted autophagy responses. Canonically, multilayered protein kinase networks converge on the conserved Atg1/ULK kinase complex (AKC) to induce non-selective and selective forms of autophagy in response to metabolic changes. Here, we show that, upon phosphate starvation, the metabolite sensor Pho81 interacts with the adaptor subunit Atg11 at the AKC via an Atg11/FIP200 interaction motif to modulate pexophagy by virtue of its conserved phospho-metabolite sensing SPX domain. Notably, we find core AKC components Atg13 and Atg17 are dispensable for phosphate starvation-induced autophagy revealing significant compositional and functional plasticity of the AKC. Our data indicate that, instead of functioning as a selective autophagy receptor, Pho81 compensates for partially inactive Atg13 during pexophagy when TORC1 remains active under phosphate starvation. Our work shows Atg11/FIP200 adaptor subunits not only bind selective autophagy receptors but also modulator subunits that convey metabolic information directly to the AKC for autophagy regulation.

## Introduction

Macroautophagy (hereafter autophagy) is a conserved catabolic process critical for cellular homeostasis and health. Cells evolved the ability to respond to a wide variety of stresses by forming double-membrane autophagosomes to deliver a remarkably broad scope of cellular substrates for vacuolar or lysosomal degradation and metabolite recycling. How cells integrate complex metabolic and functional information to tune substrate scope and timing to elicit stress-adapted autophagy responses is a key question. We are only beginning to define the metabolite, signal, and mechanistic space of autophagy regulation.

Formation of the Atg1/ULK kinase complex (AKC) marks the initial step in the hierarchical assembly of the highly conserved autophagy (Atg) protein machinery resulting in autophagosome biogenesis^1,2^. Canonically, cells translate metabolic information into activity patterns of multilayered protein kinase networks, which converge on the Atg1/ULK kinase complex (AKC) to regulate its localized assembly and activity. In budding yeast, the AKC consists of the protein kinase Atg1 and the scaffold proteins Atg13 and the Atg17 complex composed of Atg17, Atg29 and Atg31, and the selective autophagy adaptor Atg11^7–10^.

Autophagosome biogenesis can serve two distinct forms of autophagy. First, non-selective or bulk autophagy degrades a broad spectrum of substrates by stochastically encapsulating portions of the cytosol. A key step in the regulation of bulk autophagy is the phosphorylation of the core AKC component Atg13, an intrinsically disordered scaffolding protein, by TORC1^1,3,11,12^. In response to nutrient starvation, TORC1 inhibition causes dephosphorylation of Atg13 resulting in the formation of supramolecular AKC clusters defining the site of autophagosome biogenesis ^1,4,12,13^. Second, selective autophagy targets specific substrates ranging from ribosomes, protein aggregates to whole organelles including peroxisomes and mitochondria ^5,6,14^. Selective autophagy is activated, when selective autophagy receptors on the substrate surface recruit and cluster the AKC via binding to the adaptor protein Atg11 (FIP200 in mammals) to initiate autophagosome formation^15–20^. Driven by the physical interaction between the ubiquitin-like Atg8 protein, which is covalently linked to autophagic membranes, and substrate-bound receptors, nascent autophagosomes closely form around the targeted substrate excluding most other cytosolic components^6,21,22^.

Interestingly, nutrient starvation induces a composite response of bulk and selective forms of autophagy. Selective autophagy targets dysfunctional cellular components as part of the cellular quality control system, which likely contributes to the beneficial effects of autophagy during fasting for ageing and age-associated diseases^23–25^. Thus, how cells coordinate the levels of bulk and selective autophagy in terms of substrates and time in response to diverse nutrient stresses remains an open question.

Here, we investigate how metabolic signals converge on the AKC and are translated into bulk and selective autophagy in response to phosphate or nitrogen starvation ^26,27^. Strikingly, we discovered differential compositional and functional plasticity of AKC complex during the two nutrient stresses. Importantly, we identify phospho-metabolite sensor Pho81 as a modulator of selective autophagy, which directly conveys metabolic information to the AKC to regulate the turnover of peroxisomes during phosphate starvation.

## Results

### Metabolite sensor Pho81 binds the Atg1 kinase complex via Atg11

Canonically, protein kinase networks integrate metabolic information and converge on the Atg1 kinase complex (AKC) to regulate autophagy. We asked whether the induction of autophagy has similar compositional and functional requirements for the AKC in response to phosphate or nitrogen starvation. To address this question, we compared autophagic flux in wildtype (WT) cells with cells deficient for either the kinase Atg1, the selective autophagy adaptor Atg11 or scaffold proteins Atg13 and Atg17, Atg29 and Atg31 (**Fig. 1a**). To assess autophagy flux, we monitored turnover of 2GFP-Atg8, which is covalently linked to autophagic membranes and cleaved within vacuoles to generate free GFP^28^. Consistent with its essential function, the absence of Atg1 prevented any autophagic turnover in phosphate or nitrogen starvation (**Fig. 1b**). Interestingly, we detected substantial autophagy flux in Δ*atg11* cells during phosphate starvation, in contrast to a previous publication^27^. Strikingly, while both, Atg13 and Atg17 were essential for nitrogen starvation-induced autophagy consistent with previous data^1,7^, we found that Δ*atg13* and Δ*atg17* cells showed significant turnover of 2GFP-Atg8 during phosphate starvation (**Fig. 1b**). These data suggest that cells depended on Atg11 in the absence of Atg13 or Atg17 to induce autophagy in response to phosphate starvation. In support of this notion, we found that Δ*atg11*Δ*atg13* and Δ*atg11*Δ*atg17* cells were completely inhibited for autophagy during phosphate starvation (**Fig. 1b**). Strikingly, Δ*atg13*Δ*atg17*Δ*atg29*Δ*atg31* cells maintained significant 2GFP-Atg8 turnover compared with WT cells after phosphate starvation (24 h)(**Fig. 1b**), suggesting the existence of a minimal Atg1-Atg11 complex sufficient to support autophagic activity. Taken together, we find a strikingly differential functional and compositional plasticity of the AKC in response to phosphate and nitrogen starvation.

**Figure 1:**
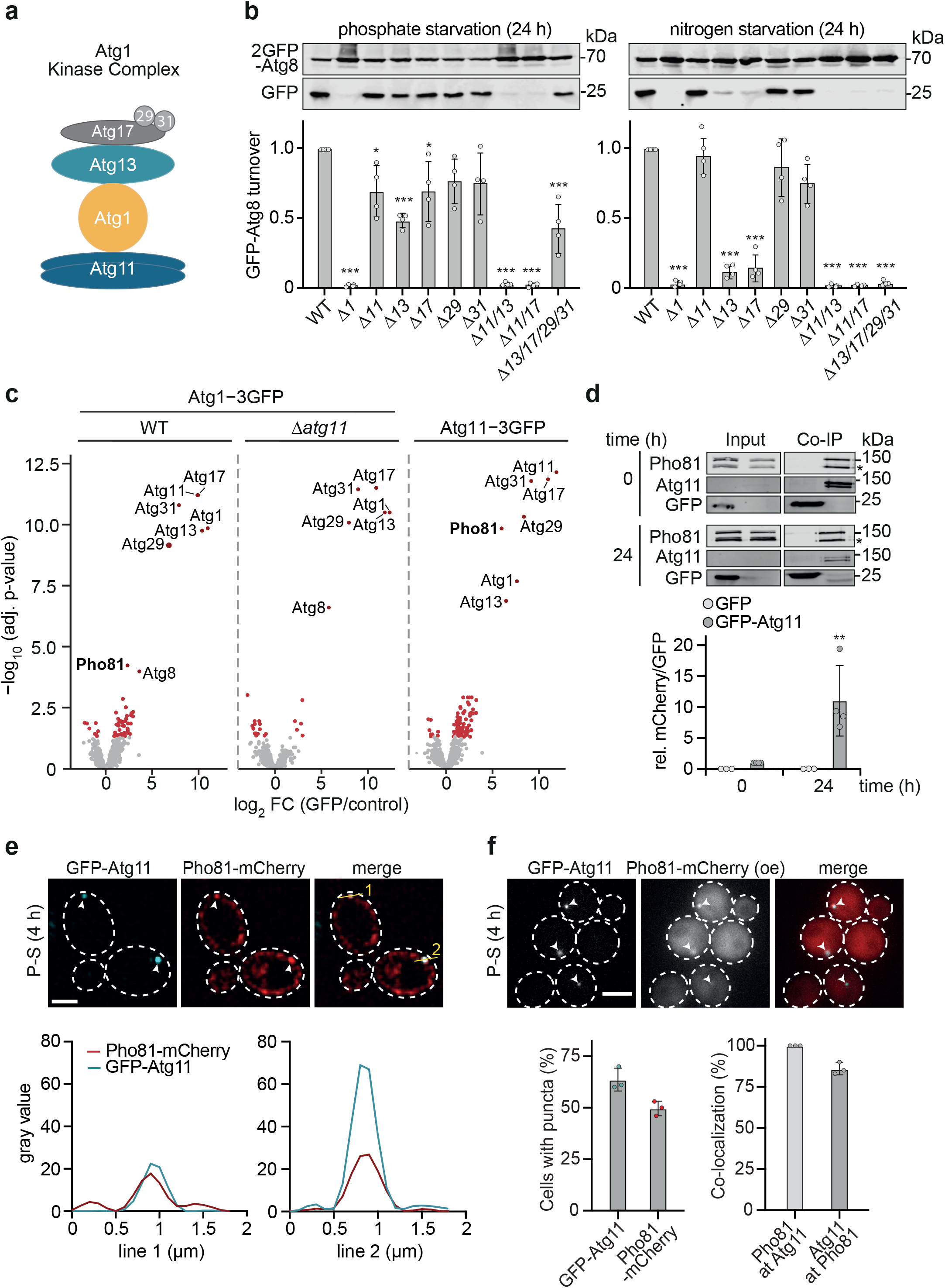
Pho81 binds to the AKC via Atg11 during phosphate starvation. **a,** Schematics of the canonical Atg1 Kinase Complex (AKC). **b,** Autophagic turnover in indicated strains expressing 2GFP-*ATG8* after P-S or N-S (24 h). Cells were analyzed by whole cell extraction and western blot analysis using an α-GFP antibody. Data are normalized to WT and are means±SD (n=4). **c**, Volcano plots of fold changes in protein abundance relative to negative control after co-immunopurification coupled with mass spectrometry (CoIP-MS) of indicated 3xGFP tagged proteins after 8 h P-S via anti-GFP microbeads. Significantly enriched proteins are shown in red (n=4). **d**, Co-immunopurification of GFP-Atg11 and Pho81-mCherry after GFP-pulldown before (0 h) and after P-S (24 h) as compared to growth condition (0 h) in strains expressing plasmid-encoded GFP-*ATG11* and *PHO81*-mCherry. Data are means ± SD (n=4). **e,** Fluorescence imaging of strains expressing pRS316-GFP-*ATG11* (cyan) and pRS315-*pho81*-mCherry (red) after P-S (4 h) and line intensity plots showing gray values of the indicated tagged protein measured along the yellow line. Scale bar is 2 μm. **f,** Fluorescence imaging of strains co-expressing plasmid encoded mCherry tagged *PHO81* under *ADH1(oe)* promoter control and GFP-*ATG11* under its endogenous promoter during growth. Scale bar is 5μm. **g,** Quantification of Pho81-mCherry and GFP-Atg11 foci per cell and the respective co-localization. Left panel: cells with respective puncta (%) and right panel: co-localization events (%), bars represent the mean ±SD. (analyzed 100 cells per image out of 3 images representing independent replicates).

Since cells displayed significant autophagy in the absence of Atg13 or the whole Atg13-17-29-31 subcomplex during phosphate starvation, we asked whether additional as yet unidentified components may be required to functionally compensate for their absence from the AKC. To address this question, we determined the protein interactome of Atg1, of Atg1 in the absence of Atg11, and of Atg11 by performing co-immunopurifications coupled with mass spectrometry (CoIP-MS) of C-terminally triple-GFP-tagged proteins isolated from cells after phosphate starvation (8 h). Interestingly, in addition to Atg8 and the canonical components of the AKC, Atg1-3GFP showed a significant enrichment for the metabolite sensor Pho81 (**Fig. 1c**). Pho81 has been shown to function as a phospho-metabolite sensor with a preference for inositol pyrophosphates and inhibitor of the cyclin-dependent Pho80-Pho85 kinase complex ^29–32^. Notably, the interaction of Pho81 with the AKC was lost when we analyzed Atg1-3GFP from Δ*atg11* cells (**Fig. 1c**), demonstrating Atg11-dependent binding of Pho81 to the AKC. Consistent with this notion, we found strong enrichment of Pho81 and all canonical AKC subunits when we profiled the protein interactome of Atg11-3GFP. Taken together, these data reveal an Atg11-dependent physical interaction of Pho81 with the AKC during phosphate starvation.

To test whether Pho81 interacts with the AKC in a phosphate starvation-dependent manner, we performed CoIP of cells co-expressing plasmid encoded GFP-*ATG11* and *PHO81*-mCherry under growing conditions or after phosphate starvation (24 h). We detected a basal interaction of Pho81-mCherry with GFP-Atg11, but not with GFP alone (**Fig. 1d**). Interestingly, the interaction drastically increased upon phosphate starvation, indicating phosphate starvation-induced binding of Pho81 to Atg11 (**Fig. 1d**). To test whether Pho81 interacts with Atg11 at sites of autophagosome formation, we analyzed cells co-expressing GFP-Atg11 with Pho81-mCherry under its endogenous promoter (**Fig. 1e**) or overexpression promoter (*ADH1*) (**Fig. 1f**). Pho81-mCherry formed punctate structures after phosphate starvation (4 h), which co-localized with GFP-Atg11 (**Fig. 1e, f**). In strains co-expressing *2GFP*-*ATG8* and plasmid encoded *PHO81*-mCherry under an *ADH1* overexpression promoter, we also observed Pho81-mCherry and GFP-Atg8 positive structures after phosphate starvation (4 h) (**Extended data 1a, b**). Taken together, our data show that Pho81 physically interacts with the AKC in an Atg11-dependent manner and localizes to Atg8- and Atg11 marked sites of autophagosome formation during phosphate starvation.

### Pho81 promotes pexophagy

We next aimed at defining the function of Pho81 at the AKC for autophagy. The function and substrates of autophagy during phosphate starvation have been unknown. To identify potential substrates of (selective) autophagy, we characterized the autophagy-dependent changes in the yeast proteome during phosphate starvation. We performed high-resolution quantitative whole cell proteomics of WT cells and Δ*atg1* cells, defective in all forms of autophagy, and Δ*atg11* cells, specifically defective in selective autophagy, after phosphate starvation (24 h). In total, we identified ~4500 proteins in a quantitative manner corresponding to > 90% of the expressed yeast proteome (**Fig. 2a, b**). During phosphate starvation, Δ*atg1* and Δ*atg11* cells significantly accumulated 263 and 294 proteins, respectively (**Fig. 2a**). Strikingly, amongst the 263 and 294 significantly enriched proteins, we identified 136 and 209 mitochondrial and 23 and 28 peroxisomal proteins for Δ*atg1* and Δ*atg11* cells compared with WT cells, respectively (**Fig. 2a, b**). These findings suggest (selective) autophagy turnover of mitochondria and peroxisomes upon phosphate starvation.

**Figure 2:**
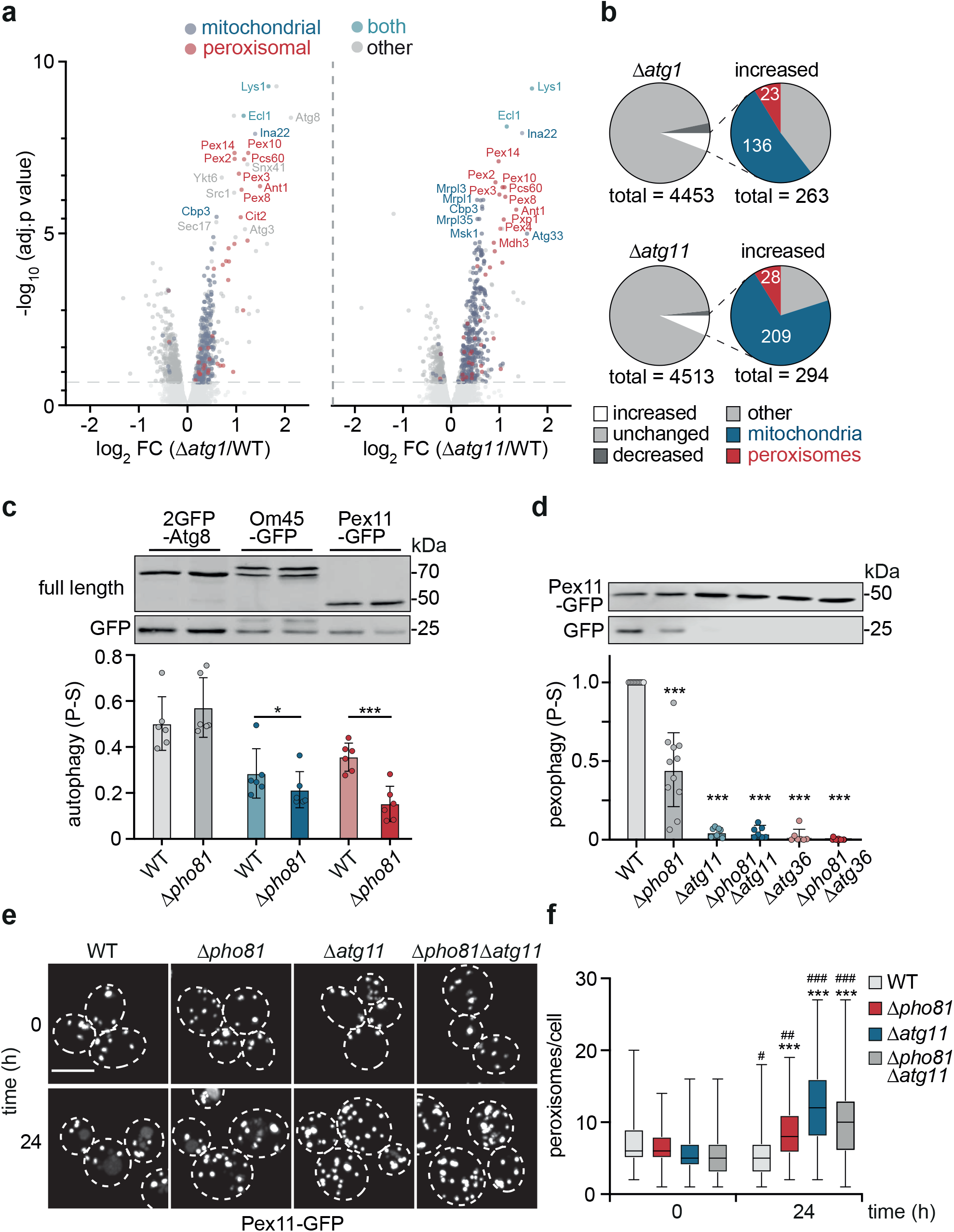
Pho81 is a positive regulator of pexophagy. **a, b** Whole cell proteome analysis using quantitative TMT labeling mass spectrometry. Proteome changes of Δ*atg1* and Δ*atg11* deletion cells as compared to WT cells after 24 h of P-S. (n=4); (**a**) Volcano plots showing relative (fold) changes; above horizontal dotted line: significantly changed as compared to WT 24 h P-S; top twenty proteins with the highest −log_10_ (adjusted p. value) are labeled; (**b**) Pie charts showing proteome changes in counts in Δ*atg1* and Δ*atg11* cells compared to WT 24 h P-S. **c**, Autophagic turnover of WT and Δ*pho81* deletion cells expressing endogenously tagged 2GFP-*ATG8*, *OM45*-*GFP* or *PEX11*-*GFP* as indicated, upon 24 h of P-S. Data shows individual data points and bars represent means ±SD (n=6). **d**, Autophagic turnover in indicated *PEX11*-*GFP* expressing deletion cells upon 24 h of P-S. Data shows individual data points and bars represent means ±SD (n=11). **e**, Images represent maximum intensity Z stack projections from 3D images of *PEX11*-GFP expressing deletion cells at 0h and 24 h of P-S. Scale bar= 5 μm. Images were deconvolved using Huygens Pro Software. **f**, Quantification of peroxisomes per cell from images shown in **e,** Boxplots show minimal to maximal value with 50% of the data points lying within the box and the black line representing the median. (Analyzed 25 cells per image out of 4 images representing independent replicates). 2-way ANOVA ***p < 0.001 comparison to WT 24 h; #p < 0.05, ##p < 0.01, ###p < 0.001 comparison over time (24h vs. 0h).

Based on these data, we asked whether Pho81 functions in mitophagy and/or pexophagy. We monitored autophagic degradation of the mitochondrial or peroxisomal membrane proteins Om45-GFP and Pex11-GFP, respectively, in comparison to 2GFP-Atg8 turnover. Pho81 did not affect autophagic membrane turnover measured by 2GFP-Atg8 consistent with previous work^27^ (**Fig. 2c**). However, in line with our proteomics data, both, mitochondria and peroxisomes were significantly degraded by autophagy during phosphate starvation (**Fig. 2c**). Interestingly, *PHO81* deletion led to a robust or mild but significant reduction of Pex11-GFP or Om45-GFP turnover after phosphate starvation (24 h), respectively (**Fig. 2c**). The selective adaptor Atg11 and the pexophagy-receptor Atg36 are essential for nitrogen starvation-induced pexophagy ^33,34^. Importantly, we found that pexophagy also depended on the presence of Atg11 and Atg36 during phosphate starvation (**Fig. 2d**). In line with autophagic turnover, the number of peroxisomes declined in WT cells after phosphate starvation (24 h) compared with non-starved cells (**Fig. 2e, f**). In contrast, reduced or blocked pexophagy in Δ*pho81*, Δ*atg11* and Δ*pho81*Δ*atg11* cells correlated with a significant accumulation of peroxisomes compared with WT cells after phosphate starvation (24 h) (**Fig. 2e, f**), establishing a physiological function for Pho81-depedent selective autophagy in the regulation of peroxisome abundance. Taken together, these data demonstrate that Pho81 promotes selective forms of autophagy with a particularly important role for pexophagy during phosphate starvation.

### Pho81 functionally interacts with Atg13 during pexophagy

The physical interaction of Pho81 with Atg11 suggests a direct role for Pho81 in selective autophagy during phosphate starvation. We tested whether Pho81 may function as a selective autophagy receptor (SAR). SARs bind both, Atg11 to cluster the AKC and Atg8 to tether selective cargos to the growing phagophore ^20^. To assess potential interactions of Pho81 with Atg8 and Atg11 using yeast-two-hybrid (Y2H), we N-terminally fused the activator domain (AD) to *PHO81* and the binding domain (BD) to *ATG11* or *ATG8*. Growth on selective plates confirmed the physical interaction of AD-Pho81 with BD-Atg11 in Y2H (**Fig. 3a**). Interestingly, we did not detect an AD-Pho81 and BD-Atg8 interaction (**Fig. 3a**). Notably, consistent with previous data^35^, the pexophagy-receptor AD-Atg36 interacted with both, BD-Atg11 and BD-Atg8, suggesting that, in contrast to Atg36, Pho81 does not function as a canonical SAR (**Fig. 3b**). In addition, the presence of Atg36 was not required for the Y2H-interaction of Pho81 with Atg11 (**Extended data 1c**). These data suggest Pho81 positively regulates Atg11-Atg36-driven pexophagy in a manner that is functionally distinct from canonical SARs.

**Figure 3:**
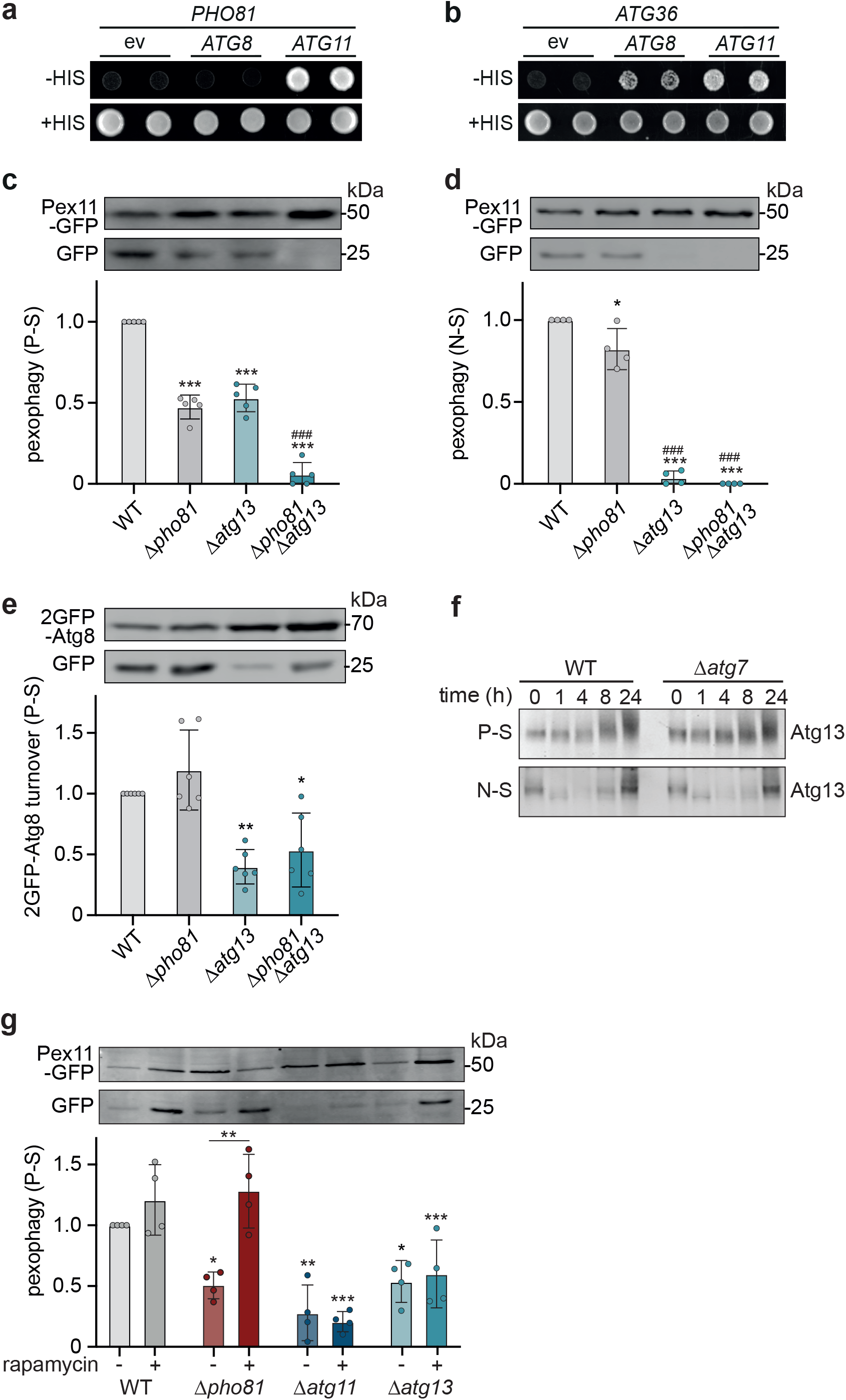
Pho81 compensates for only partially activated Atg13 during pexophagy. **a, b** Yeast-two-hybrid assay (Y2H) of cells expressing pGAD-*PHO81* or pGAD*-ATG36* in the respective combination with pGBDU-empty vector (ev), -*ATG8* or -*ATG11*. Cells were spotted on SD-complete (control) or SD-His (selection) plates and grown for 4 d. **c-e**, Autophagic turnover of (**c**) *PEX11*-*GFP* in indicated strains after 24 h P-S. (n=5) or (**d**) 24 h N-S. (n=4); (**e**) 2GFP-*ATG8* expressing strains upon 24 h P-S. (n=4). **f,** Phosphorylation of endogenous Atg13 in WT and Δ*atg7* cells during P-S and N-S at indicated timepoints. Samples were analyzed by western blot using an α-Atg13 antibody. **g,** Autophagic turnover of *Pex11*-*GFP* expressing strains after 24 h P-S ± 400ng/ml rapamycin (n=5). *p < 0.05, **p < 0.01, ***p < 0.001 compared to WT untreated or as indicated. **p < 0.01, ***p < 0.001 compared to WT or as indicated.

We observed that, in contrast to nitrogen starvation, autophagy did not strictly depend on the presence of Atg13 during phosphate starvation. Hence, we asked whether Pho81 might functionally interact with Atg13. When we assessed pexophagy, we observed a significant reduction in Pex11-GFP turnover for both, Δ*atg13* and Δ*pho81* cells compared with WT cells (**Fig. 3c**). Importantly, Δ*pho81*Δ*atg13* cells were completely impaired in pexophagy upon phosphate starvation (**Fig. 3c**), consistent with a functional interaction between Pho81 and Atg13. In contrast, while Δ*pho81* cells were only mildly affected, the presence of Atg13 was required for pexophagy during nitrogen starvation (**Fig. 3d**). Interestingly, we did not detect any additional effect on general autophagic membrane (2GFP-Atg8) turnover in Δ*pho81*Δ*atg13* cells compared with Δ*atg13* cells during phosphate starvation (**Fig. 3e**). Taken together, these data uncover a functional interaction of Pho81 and Atg13 for pexophagy during phosphate starvation.

To define the molecular mechanisms underlying the differential requirements for Atg13 and Pho81 for pexophagy during nitrogen and phosphate starvation, we examined TORC1-regulated Atg13 phosphorylation^1,4,12,13^. Consistent with previous data, TORC1 inhibition led to rapid dephosphorylation of Atg13 within 1 h in WT and Δ*atg7* cells followed by gradual re-phosphorylation during 4-24 h of nitrogen starvation (**Fig. 3f**)^1,3^. Interestingly, global phosphorylation of Atg13 was largely maintained in WT and Δ*atg7* cells upon phosphate starvation consistent with previous data (**Fig. 3f**)^27^, indicating significant TORC1 activity independent of autophagy. Given the high level of phosphorylation, we asked whether Atg13 is only partially activated and, as a consequence, cells depend on Pho81 to induce pexophagy during phosphate starvation. To address this question, we treated cells with rapamycin to inhibit TORC1 and fully activate Atg13 during phosphate starvation (**Fig. 3g**)^1^. Strikingly, Δ*pho81* cells displayed WT-like pexophagy in the presence of rapamycin (**Fig. 3g**), indicating that full Atg13 activation renders pexophagy independent of Pho81 during phosphate starvation. Importantly, pexophagy still occurred in an Atg11-dependent manner during phosphate starvation upon TORC1 inhibition, excluding the possibility of rapamycin-induced non-selective or bulk pexophagy. Interestingly, the effect of rapamycin exclusively depended on the presence of Atg13 (**Fig. 3g**), indicating that TORC1 inhibition does not affect Pho81-mediated pexophagy. In summary, our data indicate that TORC1 retains significant activity during phosphate starvation, resulting in a high level of Atg13 phosphorylation. Importantly, under these conditions, the induction of pexophagy requires the function of Pho81 to compensate for only partially activated Atg13. Taken together, our data support a model in which Pho81 is recruited to the AKC to promote pexophagy in the presence of partially inactive Atg13 during phosphate starvation. Thus, the differential compositional and functional plasticity of the AKC, we observed during phosphate and nitrogen starvation, is at least in part based on the conditional function of the modulator subunit Pho81.

### Pho81 binds Atg11 via an Atg11/FIP200 binding motif

Our data show that Pho81 binds to the AKC via Atg11 and promotes pexophagy in the presence of partially activated Atg13. We aimed at determining the molecular nature of the physical interaction of Pho81 and Atg11. Pho81 is composed of a N-terminal SPX domain, a conserved inositol (poly-/pyro-)phosphate sensor/binding domain, a minimal domain (MD), required for Pho80 binding, an ankyrin domain, and a C-terminal glycerophosphodiester-phosphodiesterase (GP-PDE) domain of unknown function (**Fig. 4a**) ^30,31^. Ankyrin domains consist of helix-turn-helix-loop repeats and commonly form a platform for protein-protein interactions defined by highly versatile loop sequences ^36^. We examined the potential role of the Pho81 ankyrin domain for Atg11 interaction using a yeast two hybrid (Y2H) approach. First, we created Pho81 variants by replacing either the entire ankyrin domain (Pho81^A^) or by systematically exchanging individual loops (Pho81^LA1^ to ^LA4^) with the corresponding sequences of the ankyrin domain of Akr1 (**Fig. 4a**). The exchange of the whole ankyrin domain (Pho81^A^ variant) impaired Pho81-Atg11 interaction, indicating a critical function for the Pho81 ankyrin domain in Atg11 binding (**Fig. 4b**). Interestingly, changes in the loop of the first Pho81 ankyrin repeat completely prevented Pho81-Atg11 interaction, but not in loops of ankyrin repeat 2-4. These data reveal a critical role for the first loop of the ankyrin domain of Pho81 for Atg11 binding.

**Figure 4:**
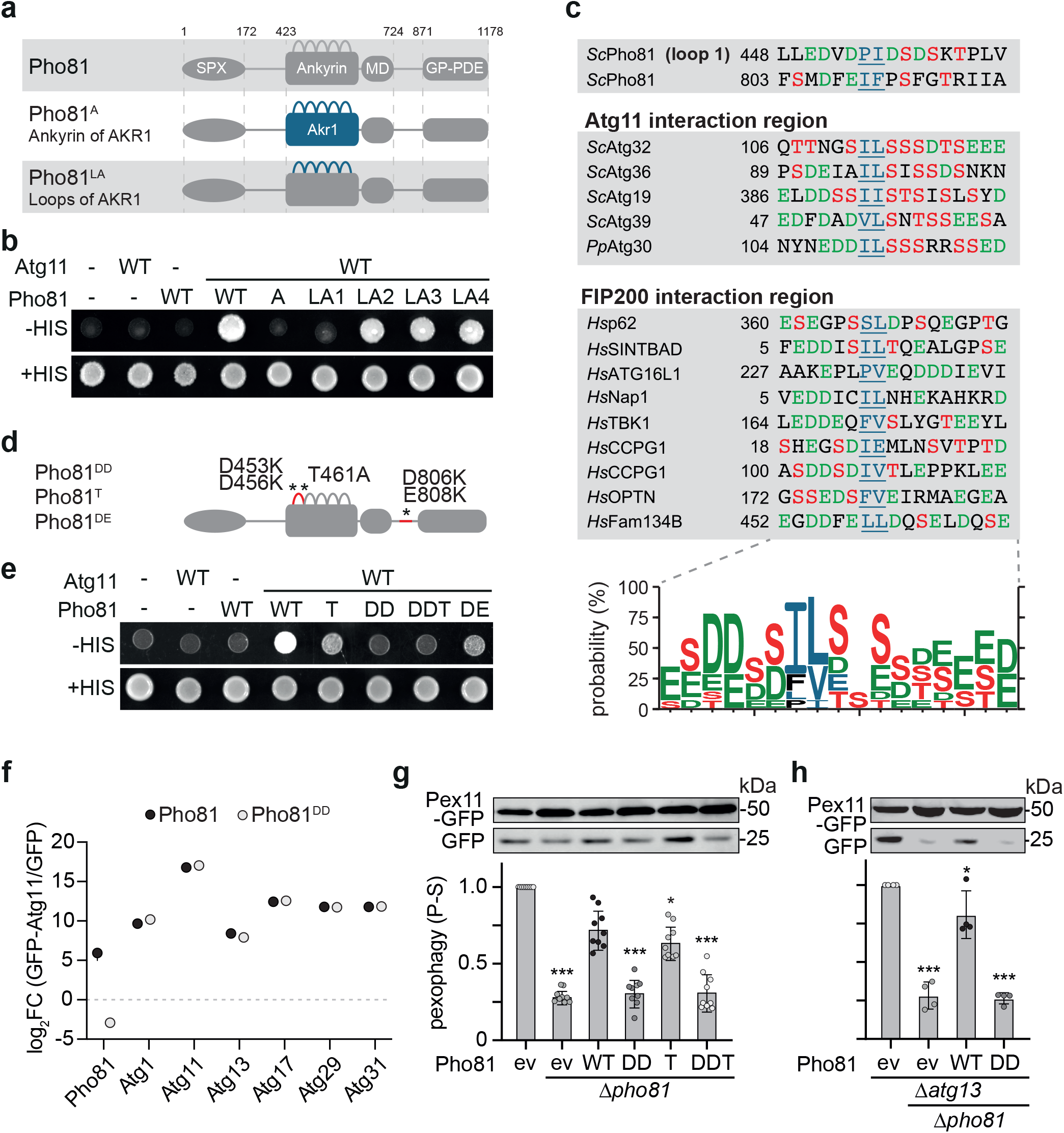
Pho81 interact with Atg11 via an Atg11/FIP200 interaction motif to promote pexophagy. **a**, Schematics of Pho81 and ankyrin repeat variants; Pho81^A^ (Pho81 ankyrin repeat replaced by Ankyrin repeat of Akr1); Pho81^LA1^ to Pho81^LA4^ (Pho81 ankyrin repeat loops 1 to 4 replaced by corresponding loop of Akr1 Ankyrin repeat loops 1 to 4). **b**, Y2H of cells expressing pGAD-*PHO81* and indicated Pho81 variants in respective combination with pGBDU-empty vector (-) or *ATG11*. 1 OD_600_ were spotted on SD-complete (control) or SD-His (selection) agar plates and grown for 4 days. **c**, Putative consensus motif of Atg11/FIP200 interaction region. The homology logo was created using WebLogo 3.0 ^37^ upon aligning Atg11/FIP200 interaction regions represented and shows the probability (%) to find a certain amino acid at the respective position. *Sc*, Saccharomyces cerevisiae; *Pp*, Pichia pastoris; *Hs*, Homo sapiens**. d**, Schematics of the mutations introduced in the Pho81^DD^, Pho81^T^ and Pho81^DE^ variants, based on the Atg11/FIP200 binding region analysis in (**c). e**, Y2H of cells expressing pGAD-*PHO81* and indicated Pho81 variants in respective combination with pGBDU-empty vector (-) or *ATG11*. Cells were spotted on SD-complete (+HIS) or SD-His (-HIS) agar plates and grown for 4 days. **f**, Log_2_ FC enrichment of proteins bound to plasmid encoded GFP-*ATG11* and detected by CoIP-MS using anti-GFP microbeads from cells co-expressing plasmid encoded *PHO81*-mCherry or *pho81*^DD^-mCherry. CTRL is GFP (n=4). Statistical analysis is shown in table S2. **g**, **h,** Autophagic turnover of cells expressing plasmid encoded Pho81 variants as indicated in the respective *PEX11-GFP* expressing deletion cells upon 24 h of P-S. Data show individual data points relative to WT. Bars are means ±SD ((**g**) n=9; (**h**) n=4).

To test our findings from the Y2H assay, we co-expressed plasmid-encoded *PHO81*-mCherry under the *ADH1*-promoter with GFP-*ATG11* under its endogenous promoter and monitored co-localization during growing conditions using fluorescence microscopy. We observed quantitative recruitment of Pho81-mCherry to GFP-Atg11 foci, establishing Pho81-mCherry puncta formation as a proxy for Atg11 interaction (**Fig. 1f**). Consistent with our Y2H data, foci formation specifically required Pho81 ankyrin repeat loop 1, but not loop 2 and loop 3 (**Extended Data 1d, e**). Interestingly, the Pho81^LA4^ variant allowed Atg11 interaction in Y2H, but failed to form foci likely due to the observed nuclear localization (**Fig. 4b, Extended Data 1d, e**). In summary, these data demonstrate that Pho81 binding and recruitment to Atg11-marked sites of autophagosome biogenesis specifically depend on the loop of the first ankyrin repeat of Pho81.

We aligned known protein sequences required for Atg11- and FIP200-binding to identify common features of Atg11/FIP200 interaction regions (FIRs) (**Fig. 4c**)^18,37–39^. We found that FIRs display a cluster of negatively charged aspartic and glutamic acid (D or E) or phosphorylatable serine or threonine (S or T) residues proximal to two hydrophobic amino acids (P, V, L, I or F) (**Fig. 4c**), consistent with previous data^18,39^. Interestingly, we identified potential Atg11 interaction motifs in the loop of the first Pho81 ankyrin repeat (aa 451-461) and at aa 806-816 located between the minimal and GP-PDE domain (**Fig. 4c**). To test the functional relevance of these predicted FIR motifs for Atg11 binding, we generated Pho81^DD^ (D453K/D456K), Pho81^T^ (T461A), Pho81^DDT^ (D453K/D456K/T461A) and Pho81^DE^ (D806K/E808K) variants by mutating the corresponding residues to lysine (K) or alanine (A) as indicated (**Fig. 4d**). Interestingly, consistent with the presence of a FIR, Y2H analysis revealed a strict requirement for D453/D456 in loop 1 (Pho81^DD^) for the Pho81-Atg11 interaction. Additionally, we found a partial reduction in Atg11 binding for the Pho81^DE^ variant (**Fig. 4e**). The residue T461 in Pho81 is a confirmed phosphorylation site, which is at a similar position within the FIR motif as the phosphorylatable S119 site in the yeast mitophagy receptor, Atg32 ^35,40–42^. Phosphorylation of S119 supports the interaction of Atg32 with Atg11 ^35,40–42^. The Pho81^T^ variant had a partial Atg11 binding defect in the Y2H assay. Taken together, our data suggest T461 and D806/E808 sites are not essential but support the Pho81-Atg11 interaction (**Fig. 4d, e**). Critically, CoIP-MS of plasmid encoded GFP-*ATG11* from cells expressing either *PHO81-* or *pho81*^*DD*^-*mCherry* validated the D453/D456-dependent interaction of Pho81 and Atg11 after phosphate starvation (8 h) (**Fig. 4f**). Notably, the interaction of the core AKC components with GFP-Atg11 was unchanged in the presence of Pho81 or Pho81^DD^, demonstrating that the core AKC assembles in a manner that is independent of Pho81-Atg11 interaction (**Fig. 4f**). Our data identify an Atg11/FIP200 interaction motif within the first repeat of the ankyrin domain of Pho81 required for Atg11-dependent binding to the AKC.

The identification of an Atg11-binding-deficient Pho81 variant allowed us to specifically assess the functional relevance of the interaction of Pho81 with the AKC for pexophagy. Expression of plasmid-encoded *PHO81* under its endogenous promoter restored pexophagy in *Δpho81* cells, while *pho81^DD^* and *pho81^DDT^* did not (**Fig. 4 g**). Expression of *pho81^T^* slightly reduced pexophagy levels, suggesting that posttranslational modifications of Pho81 at T461 may contribute to regulate Pho81-dependent pexophagy (**Fig. 4g**) ^35,39,42^. Notably, impaired Pho81-Atg11 interaction through mutation of D453/D456 strongly blocked pexophagy in the absence of Atg13 (**Fig. 4h**), indicating Pho81 binding to Atg11 is required for its compensatory function for pexophagy. To critically test whether the effects of the Pho81 variants on pexophagy were caused by potential changes in the PHO pathway, we analyzed pexophagy in the absence of the transcription factor *PHO4*, essential for PHO pathway activation ^29,43^. In contrast to cells expressing *PHO81*, the *pho81*^DD^ variant failed to restore pexophagy in Δ*pho81*Δ*pho4* cells (**Extended data 1f**). Taken together, our data show that the binding of Pho81 to Atg11 via an Atg11/FIP200 interaction motif at the AKC is required for Pho81-mediated pexophagy. Importantly, independent of its role in the PHO pathway, Pho81 functions as modulator subunit of the AKC without affecting core subunit assembly.

### The metabolite sensing SPX domain of Pho81 promotes pexophagy

SPX domains are evolutionarily conserved sensors of phospho-metabolites from yeast to humans ^29,31,44^. We hypothesized that the SPX domain of Pho81 senses phospho-metabolites to regulate pexophagy at the AKC during phosphate starvation. SPX domains are small (135-380 aa) N-terminal structures with two long core helices, which preferentially interact with inositol polyphosphates IP_6_ and pyrophosphates IP_7_ via a highly conserved cluster of amino acid residues ^31,45^. This phosphate binding cluster maps onto Y24, K28 and K154 in Pho81 (**Fig. 5a, b**) ^31^. We generated a Pho81^ΔSPX^ variant without SPX domain (1-172 aa) or a Pho81^YKK^ variant with mutated residues Y24A, K28A and K154A, predicted to be deficient for phospho-metabolite binding (**Fig. 5a, b**)^31^. CoIP-MS analysis of GFP-Atg11 demonstrated that changes in the SPX domain did not affect the interaction of neither Pho81 with the AKC nor the AKC components with each other (**Fig. 5c**). Importantly, the presence of the SPX domain and its functional integrity were required for Pho81 to compensate for the absence of Atg13 during phosphate starvation-induced pexophagy (**Fig. 5d**). Metabolite sensing by the SPX domain of Pho81 regulated pexophagy in a manner that was independent of the PHO pathway (**Extended data 1f**). Taken together, these data strongly support a critical role for the phospho-metabolite sensing function of the SPX domain of Pho81 for pexophagy during phosphate starvation.

**Figure 5:**
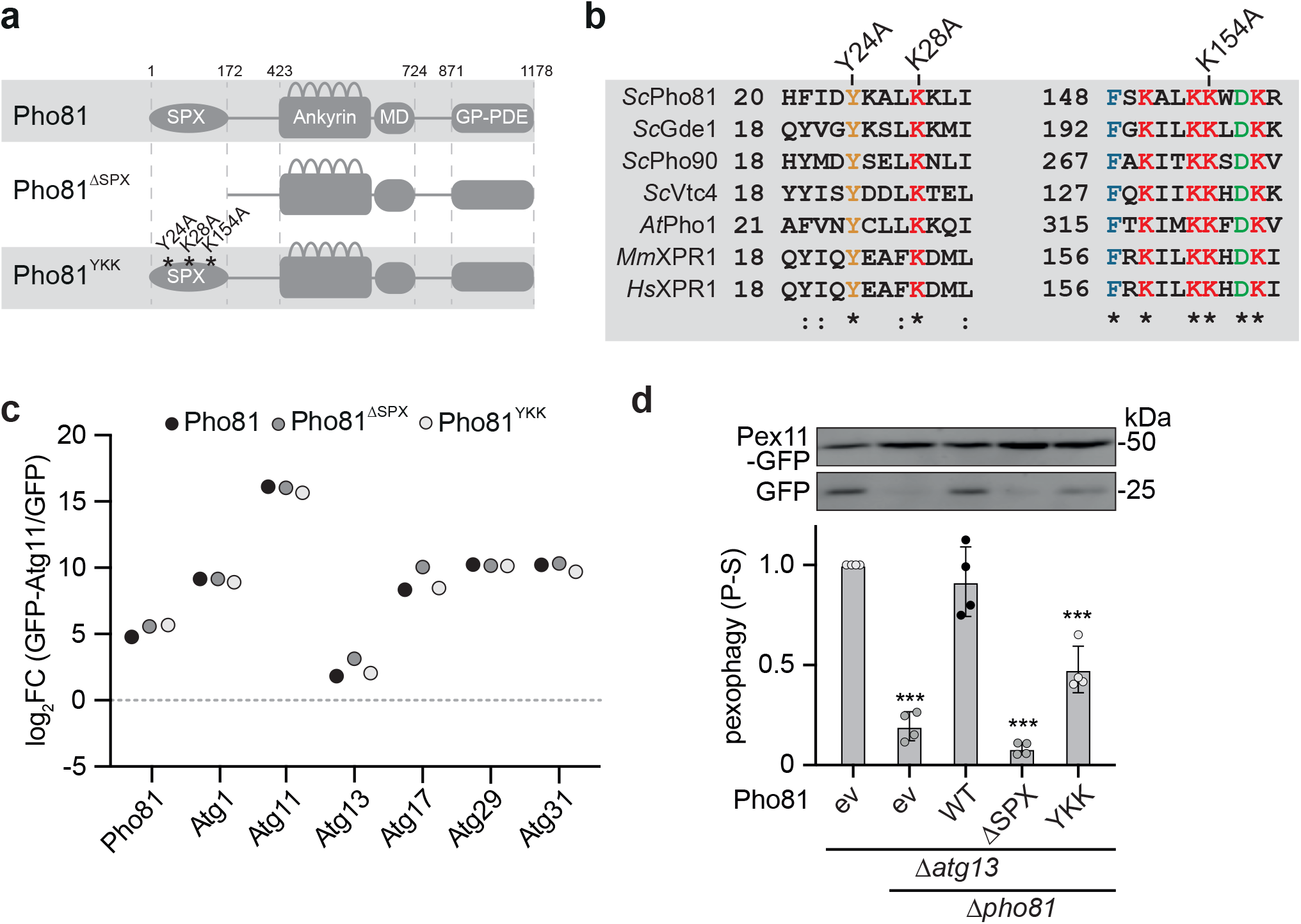
The inositol phosphate sensing SPX domain of Pho81 regulates pexophagy. **a**, Schematics of the Pho81 SPX domain variants; Pho81^ΔSPX^ (C-terminus 173-1178); Pho81^YKK^ (alanine exchange mutant of Y24, K28 and K154, the residues conserved for inositol phosphate sensing^31^). **b,** Sequence alignment of SPX domain containing proteins from *Sc*, Saccharomyces cerevisiae; *At*, Arabidopsis thaliana; *Mm*, Mus musculus; *Hs*, Homo sapiens; as indicated using ClustalO. **c**, Log_2_ FC enrichment of GFP-Atg11 bound proteins detected by CoIP-MS using anti-GFP microbeads from cells co-expressing plasmid encoded mCherry tagged *PHO81*, *pho81*^YKK^ and *pho81*^ΔSPX^ variants. CTRL is GFP (n=4). Statistical analysis is shown in table S2. **d,** Autophagic turnover of cells expressing indicated plasmid encoded Pho81 variants in the respective *PEX11*-GFP expressing deletion cells upon 24 h of P-S. Data shows individual data points relative to WT 24h and bars represent means ±SD (n=4).

## Discussion

How cells convert metabolic information into stress-adapted autophagy responses is a key question. Our work describes a molecular mechanism, which conveys metabolic information directly to the AKC largely independent of upstream protein kinase networks. Specifically, by comparing phosphate and nitrogen starvation in budding yeast, our work uncovers significant compositional and functional plasticity of the AKC in dependence of metabolic stress. In parts, this plasticity is based on the presence of a conditional subunit of the AKC, Pho81. Pho81 binds to Atg11 via an Atg11/FIP200 interaction motif at the AKC to regulate pexophagy upon phosphate starvation. Interestingly, our work indicates that, rather than functioning as a canonical autophagy receptor in parallel with Atg36, Pho81 promotes pexophagy by compensating for partially inactive Atg13 due to significant activity of TORC1 during phosphate starvation. Upon nitrogen starvation-induced TORC1 inhibition, the intrinsically disordered protein Atg13 cooperates with the Atg17 complex in organizing supramolecular AKC clusters^12,19,46^. Thus, we hypothesize that Pho81 aids in AKC clustering when Atg13 is only partially dephosphorylated during phosphate starvation. Notably, our data suggest the phospho-metabolite sensing function of the conserved SPX domain is required for Pho81 to compensate for the absence of Atg13 and to drive pexophagy. Our work supports a model in which Pho81 binds to the adaptor protein Atg11 as modulator subunit to convey metabolic information directly to the AKC to regulate selective autophagy.

Based on our work, we propose that Atg11/FIP200 subunits bind two classes of proteins, which can be conceptually categorized into receptor and modulator proteins. Excitingly, our and work from others suggests the existence of a diverse class of modulator proteins of the AKC. For example, the induction of xenophagy upon bacterial invasion in mammalian cells requires the association of the protein kinase TBK1 with FIP200, which is mediated by two adaptor proteins, NAP1/SINTBAD, containing FIP200 binding motifs^37^. TNIP1 negatively regulates mammalian mitophagy in parts by binding to FIP200 via a potential FIP200 interaction motif^47^. FIP200 also binds Atg16L1 via a FIP200 interaction motif to promote starvation-induced autophagy in mammalian cells^48,49^. These few examples already demonstrate the remarkable versatility of modulator proteins in modifying the activity and functions of the core AKC. We are likely only beginning to explore the scope of potential modulator subunits of Atg11/FIP200.

Excitingly, our mechanistic work has the potential to translate into therapeutical approaches, because Pho81 is essential for fungal infection ^44^. In a mouse model, *Cryptococcus neoformans* expressing Pho81 variants with defects in their SPX domain fail to establish lung and brain infections due to their inability to sense inositol pyrophosphates ^44^. In line with our finding that Pho81 functions in autophagy independent of its role in the PHO pathway in budding yeast, virulence is only partially reduced upon inhibition of the PHO pathway in *Cryptococcus neoformans* ^50^. Thus, our work on the function of Pho81 as a metabolite sensing modulator of selective autophagy may provide mechanistic insights with the potential for the development of novel antifungal drugs ^44,51^.

## Acknowledgments

We would like to thank Lena Pernas, Sebastian Hofer and all members of the Graef group for discussion; Prof. Daniel Klionsky for the α-Atg13 primary antibody; the members of the Proteomics Core and FACS & Imaging Facility at the Max Planck Institute for Biology for excellent support. This work was supported by the Max Planck Society and the Deutsche Forschungsgemeinschaft (DFG-German Research Foundation) - SFB 1218 - project number 269925409 to M. Graef.

## Author contributions

A.G., R.G. and M.G. conceptualized the study; A.G., R.G., M.S., E.R., A.R. and M.G designed and performed experiments and analyzed and interpreted the data; A.G. and M.G. wrote the manuscript. All authors approved the final version of the manuscript.

## Abbreviations

P-S: phosphate starvation
N-S: nitrogen starvation
AKC: Atg1 kinase Complex
SAR: selective autophagy receptor
FIR: Atg11/Fip200 interaction region
Y2H: yeast two hybrid
AD: activator domain
BD: binding domain
Co-IP-MS: Co-Immunoprecipitation coupled with mass spectrometry

## Figure Legends

**Extended Data 1**

**a,** Fluorescence imaging of strains co-expressing plasmid encoded mCherry tagged *PHO81* under an *ADH1* promoter and endogenously tagged 2GFP-*ATG*8 upon 4 h P-S. Scale bar= 5μm. **b,** Quantification of Pho81-mCherry puncta formation and co-localization with 2GFP-Atg8 positive structures; each dot represents the total of puncta in 100 cells ±SD. **c,** Y2H of cells expressing pGAD-empty vector (ev) and *PHO81* and pGBDU-*ATG11* in WT (pJ69-4a) cells and Δ*atg36* deletion cells. 1 OD_600_ were spotted on SD-complete (control) or SD-His (selection) agar plates and grown for 4 days. **d,** Fluorescence imaging of strains expressing plasmid encoded mCherry tagged Pho81 variants under *ADH1* promoter control during growth. Scale bar= 5μm. **e,** Quantification of Pho81-mCherry puncta per cell; each dot represents the number of cells with Pho81-mCherry foci in (%) ±SD. (analyzed ≥ 25 cells per image out of 5 images representing independent replicates)**. f,** Autophagic turnover of cells expressing indicated plasmid encoded Pho81 variants in the respective *PEX11*-GFP expressing deletion cells upon 24h of P-S. Data shows individual data points relative to WT 24h and bars represent means ±SD (n=6).

## Materials and Methods

### Yeast Strains and Media

The strains used in this study were derivates of W303. For Yeast two hybrid we used derivates of PJ69-4a. All strains are listed in Table S1. Gene deletions were generated by replacing the complete ORF with indicated marker cassettes using PCR-based targeted homologous recombination as described (Sheff and Thorn, 2004). Plasmids were constructed by in vivo “gap repair” based on homologous recombination (Oldenburg et al. 1997) into HindIII- and EcoRI-linearized pRS315 and pRS316 or BamHI and EcoRI-linearized pGAD and pGBDU, respectively. To modify genes to express functional C-terminal tags, the following templates were used pFA6a-*link*-*yEGFP*-*CaURA3*, pFA6a-*link*-*3yEGFP*-*CaURA3*, pFA6a-*link*-*mCherry*-*kanMX6*, pFA6a-*link*-*mCherry*-*HIS3* as previously described (Graef et al. 2013, Morawska and Ulrich, 2013). Strains were grown at 30°C in synthetic complete medium SDC (0.7% (w/v) yeast nitrogen base (BD Difco) and 2% (w/v) α-D-glucose (Sigma)) or synthetic medium lacking respective amino acids for plasmid selection. For starvation experiments, cells were harvested from early log-phase, washed 4 times and resuspended in the respective starvation medium. For phosphate starvation in SD-Pi (0.15% (w/v) yeast nitrogen base without amino acids, ammonium sulphate and phosphate, supplemented with KCl (Foremedium), 2% (w/v) α-D-glucose (Sigma), 0.5% (w/v) ammonium sulfate (Sigma)). For nitrogen starvation in SD-N medium (0.17% (w/v) yeast nitrogen base without amino acids and ammonium sulfate (BD Difco) and 2% (w/v) α-D-glucose (Sigma)). For rapamycin treatment stock solution (400 μg/ml in DMSO; Calbiochem #553210) was diluted 1:1000 to a final concentration of 400 ng/ml, CTRL cells were mock treated with equal volume of DMSO.

### Yeast two Hybrid Assay

For the Yeast two Hybrid Assay 1 OD600-units of the respective strain were serially diluted (1:10) and four concentrations were spotted on the respective agar selection plates (SD medium containing 2% (w/v) agar and the defined amino acid composition). Growth was documented by plate imaging using the Biorad Chemidoc IP Imaging System.

### Fluorescence microscopy and image analysis

Cells were transferred to 96-well glass bottom microplates (Greiner Bio-One) containing indicated media. For fixed time point analyses, cells were imaged at room temperature with an inverted microscope (Nikon Ti-E) using a Plan Apochromat IR 60x 1.27 NA objective (Nikon) and Spectra X LED light source (Lumencor). Three-dimensional light microscopy data were collected using the triggered Z-Piezo system (Nikon) and orca flash 4.0 camera (Hamamatsu). High-speed confocal imaging was performed with a Dragonfly 500 series spinning disk microscope (Andor, Oxford instruments) equipped with a Zyla 4.2 Plus sCMos camera (Andor) using a Lamba CFI-Plan Apochromat 60x 1.4 NA oil immersion objective (Nikon). Three-dimensional data were processed using Fusion software (Andor), and Fiji ImageJ Version 2.1.0. Deconvolution was performed using Huygens Professional 16.10 where indicated. Data are shown as single sections if not stated otherwise.

### Whole cell extraction, western blot analysis and quantification

Cells corresponding to 0.4 OD_600_-units were collected and lysed by alkaline whole cell extraction using 0.255 M NaOH (Roth). Proteins were precipitated with 50% (w/v) trichloroacetic acid (Roth) and washed once with ice-cold acetone (Merck). Protein pellets were resuspended in 1 x sodium dodecyl sulphate (SDS) sample buffer (50 mM Tris/HCl pH 6.8 (Roth), 10% (v/v) glycerol (Sigma), 1% (w/v) SDS (Roth), 0.01% (w/v) bromphenolblue (Roth), 1% (v/v) β-mercaptoethanol (Merck)).

Protein extracts were analyzed by SDS-PAGE and immunoblotting using α-GFP (monoclonal, Takara), α-mCherry (polyclonal, Genetex), α-Atg13 (polyclonal, a gift from Prof. Daniel Klionsky) antibodies. Primary antibodies were visualized using corresponding secondary α-mouse or α-rabbit Dylight800/680 antibodies (Rockland Immunochemicals) and signal intensities were quantified using the Li-COR Odyssey Infrared Imaging system (Biosciences).

### Whole cell extraction for mass spectrometry analysis

Cells corresponding to 1 OD_600_-units were collected in 2.0 ml safe lock tubes (Eppendorf 0030 121.880), washed 1x with H_2_O and snap frozen in liquid nitrogen. Cells were disrupted using Tissue Lyser (Qiagen) 1min 25 Hz. 1ml −20°C cold Extraction Buffer MTBE::MeOH 75::25 (v/v) (MTBE (Sigma 306975); MeOH (Biosolve 136841)) was added and samples were vortexed thoroughly, ultrasonicated 10min in bath-type sonicator cooled with ice and incubated on a thermomixer at 4°C for 30 min, centrifuged 10 min at 4°C and 16.000 g. Supernatant was removed and pellet (containing proteins) dried under the hood (and if necessary stored at −80°C until further use). The pellet was resolved in 50 μl Urea (8 M) and 0.4 μl TCEP (0.25 M) and 0.69 μl CAA (0.8 M) were added prior 1h incubation at room temperature. 1 μl Lys-C (0.5 μg/μl) (Endoproteinase MS Grade, 90051, Life Technologies) were added and incubated for min. 2h at room temperature. 150 μl Ammonium Bicarbonate (50mM) (Sigma) were added prior trypsinization with 1 μl trypsin (1 μg/μl) (Trypsin-gold V5280 Promega) over night at 37°C. Reactions were stopped by the addition of 0.1% (v/v) trifluoroacetic acid (Sigma).

### Co-immunoprecipitations

Cells corresponding to 125 OD_600_-units (for subsequent Western Blot Analysis) or 250 OD_600_-units (for Mass Spectrometry coupled analysis) were pelleted, washed and resuspended in lysis buffer (50 mM Tris pH 7.5; 150 mM NaCl; 1 mM MgCl_2_; 10% Glycerol; Complete protease inhibitor (Roche); 1 mM PMSF or 2 mM AEBSF-hydrochloride (BioChemica APA1421.0001). Cell pellets were created in liquid N_2_ and stored at −80°C until further processing.

Frozen cell pellets were lysed using a Freezer/Mill (6970EFM, SPEX^®^ Metuchen, NJ; settings T1 = 5 min; T2 = 2 min; T3 = 2 min; cycles, 2; rate, 7). Thawed cell lysates were cleared by two consecutive centrifugation steps at 3000 rpm and 12000 rpm at 4°C for 10 min each (Heraeus Multifuge X3R, TX-1000 rotor, Thermo Scientific). After clearance 0.25% “NP-40 alternative” (Milipore 492016) were added to supernatants. Supernatants were incubated with μMACS anti-GFP Microbeads (Miltenyi Biotec) for 1h at 4°C. Beads were isolated using Miltenyi μ-columns and a MultiMACS^®^ M96 Seperator (Miltenyi Biotec). Columns were equilibrated with lysis buffer, loaded and washed 2x with washing buffer 1 (50 mM Tris pH 7.5; 150 mM NaCl; 10% Glycerol; 0.25% “NP-40 alternative”) and 2x with washing buffer 2 (50 mM Tris pH 7.5; 150 mM NaCl; 10% Glycerol)

For analysis by western blotting, samples were eluted by addition of 100 μL SDS-sample buffer. For analyses by mass spectrometry, on-bead digestion was performed by addition of 100 μL trypsinization buffer (50 mM ammonium bicarbonate, Sigma and 300 ng trypsin per column, Trypsin-gold V5280 Promega) for 1 h at 23°C. Samples were eluted by addition of 150 μL 50 mM ammonium bicarbonate. After overnight incubation at 23°C, reactions were stopped by the addition of 0.1% (v/v) trifluoroacetic acid (Sigma). The aqueous solution was evaporated in a SpeedVac (Eppendorf) and analyzed by *Nano*-*ESI*-*MS*/*MS analysis or high*-*resolution mass spectrometry*.

### Proteomics analysis

Peptides were rescued according Li et al.^53^ separated on a 25 cm, 75 μm internal diameter PicoFrit analytical column (New Objective) packed with 1.9 μm ReproSil-Pur 120 C18-AQ media (Dr. Maisch,) using an EASY-nLC 1200 (Thermo Fisher Scientific). The column was maintained at 50°C. Buffer A and B were 0.1% formic acid in water and 0.1% formic acid in 80% acetonitrile. Peptides were separated on a segmented gradient from 6% to 31% buffer B for 40 or 80 min and from 31% to 50% buffer B for 5 or 10 min at 200 nl/min. Eluting peptides were analyzed on a QExactive HF mass spectrometer (Thermo Fisher Scientific). Peptide precursor m/z measurements were carried out at 60000 or 12000 resolution in the 300 to 1800 m/z range. The top ten most intense precursors with charge state from 2 to 7 only were selected for HCD fragmentation using 25% normalized collision energy. The m/z values of the peptide fragments were measured at a resolution of 30000 using a minimum AGC target of 8e^3^ and 80 ms maximum injection time. Upon fragmentation, precursors were put on a dynamic exclusion list for 20 or 45 sec.

For the analysis of the total proteome, four micrograms of desalted peptides were labeled with tandem mass tags (TMT10plex, Thermo Fisher Scientific cat. No 90110) using a 1:20 ratio of peptides to TMT reagent. TMT labeling was carried out according to manufacturer’s instruction with the following changes: dried peptides were reconstituted in 9 μL 0.1M TEAB to which 7 μL TMT reagent in ACN was added to a final ACN concentration of 43.75%, the reaction was quenched with 2 μL 5% hydroxylamine. Labeled peptides were pooled, dried, resuspended in 0.1% formic acid (FA), split into two samples, and desalted using home-made STAGE tips. One of the two samples was fractionated on a 1 mm x 150 mm ACQUITY column, packed with 130 Å, 1.7 μm C18 particles (Waters cat. no SKU: 186006935), using an Ultimate 3000 UHPLC (Thermo Fisher Scientific). Peptides were separated at a flow of 30 μL/min with a 96 min segmented gradient from 1% to 50% buffer B for 85 min and from 50% to 95% buffer B for 11 min; buffer A was 5% ACN, 10mM ammonium bicarbonate (ABC), buffer B was 80% ACN, 10 mM ABC. Fractions were collected every three minutes, and fractions were pooled in two passed (1 + 17, 2 + 18 … etc.) and dried in a vacuum centrifuge (Eppendorf). Dried fractions were resuspended in 0.1% formic acid and analyzed on a Orbitrap Lumos Tribrid mass spectrometer (Thermo Fisher Scientific) equipped with a FAIMS device (Thermo Fisher Scientific) that was operated in two compensation voltages, −50 and −70. Synchronous precursor selection based MS3 was used for TMT reporter ion signal measurements. Raw files were split based on the FAIMS compensation voltage using FreeStyle (Thermo Fisher Scientific).

### Protein identification and quantification

The raw data were analyzed with MaxQuant version 1.5.3.17 or version 1.6.0.16 ^54^ using the integrated Andromeda search engine ^55^. The total proteome data was analyzed using MaxQuant version 1.6.10.43. Peptide fragmentation spectra were searched against the yeast proteome (downloaded May 2017, February 2018 or October 2018 from UniProt). Methionine oxidation and protein N-terminal acetylation were set as variable modifications; cysteine carbamidomethylation was set as fixed modification. The digestion parameters were set to “specific” and “Trypsin/P,” The minimum number of peptides and razor peptides for protein identification was 1; the minimum number of unique peptides was 0. Protein identification was performed at a peptide spectrum matches and protein false discovery rate of 0.01. For the analysis of the total proteome samples, the isotope purity correction factors, provided by the manufacturer, were included in the analysis. The “second peptide” option was on. Successful identifications were transferred between the different raw files using the “Match between runs” option. Label-free quantification (LFQ)^56^ was performed using an LFQ minimum ratio count of 2. LFQ intensities were filtered for at least four valid values in at least one group and imputed from a normal distribution with a width of 0.3 and down shift of 1.8. Differential expression analysis was performed using limma ^57^.

### Quantification and Statistical Analysis

Error bars represent SD as indicated in the figure legends. Box and whiskers plots: the boxes extend from the 25^th^ to the 75^th^ percentile spanning the inter-quartile range (IQR), the whiskers show the minimal and the maximal value (min to max), the line in the middle of the boxes represents the median. Data were processed in Graphpad Prism 8.

Statistical analysis was performed using an ordinary one-way ANOVA with Tukey’s multiple comparisons test of one independent variable or a two-way ANOVA with Sidak’s multiple comparisons test of two independent variables, as indicated. *, p < 0.05; **, p < 0.01; ***, p < 0.001. Figures and graphs were assembled in Adobe Illustrator Version 24.3.

