## Supplementary material for "A metabolite sensor subunit of the Atg1/ULK complex regulates selective autophagy": Table S1

Table S1: Strains used in this study

| Position | Genotype | |
|---|---|---|
| YMG1 | W303 *ade2-1; leu2-3; his3-11,15; trp1-1; ura3-1; can1-100* | (Thomas and Rothstein 1989) |
| YMG8 | W303 pRS306-*prATG8-2xyeGFP-ATG8* | (Velasquez et al., 2016) |
| AG 4 | W303 *Δatg1::natMx6* pRS306-*prATG8-2xyeGFP-ATG8* | This manuscript |
| YMG9 | W303 *Δatg7::kanMx6* pRS306-*prATG8-2xyeGFP-ATG8* | (Velasquez et al., 2016) |
| AG 6 | W303 *Δatg11::natMx6* pRS306-*prATG8-2xyeGFP-ATG8* | This manuscript |
| AG 7 | W303 *Δatg13::natMx6* pRS306-*prATG8-2xyeGFP-ATG8* | This manuscript |
| AG 8 | W303 *Δatg11::natMx6 Δatg13::kanMx6* pRS306-*prATG8-2xyeGFP-ATG8* | This manuscript |
| AG 203 | W303 *Δatg17::natMx6* pRS306-*prATG8-2xyeGFP-ATG8* | This manuscript |
| AG 204 | W303 *Δatg11::natMx6 Δatg17::KanMx6* pRS306-*prATG8-2xyeGFP-ATG8* | This manuscript |
| AG 205 | W303 *Δatg29::natMx6* pRS306-*prATG8-2xyeGFP-ATG8* | This manuscript |
| AG 206 | W303 *Δatg31::natMx6* pRS306-*prATG8-2xyeGFP-ATG8* | This manuscript |
| AG 207 | W303 *Δatg13::natMx6 Δatg17::kanMx6 Δatg29::hphMx6 Δatg31::his3Mx6* pRS306-*prATG8-2xyeGFP-ATG8* | This manuscript |
| AG 257 | W303 *Δatg19::HIS3 ATG1-3xyeGFP-caURA3* | This manuscript |
| AG 283 | W303 *Δatg19::HIS3 Δatg11::natmx6 ATG1-3xyeGFP-caURA3* | This manuscript |
| AG 250 | W303 *Δatg19::HIS3 ATG11-3xyeGFP-caURA3* | This manuscript |
| AG 862 | W303 *Δpho81::kanMx6* pRS316-*prATG11- yeGFP* -ATG11-*hphMX6* pRS315-*prPHO81-PHO81-mCherry-his3mx6 ade2::ADE2* | This manuscript |
| AG 864 | W303 *Δpho81::kanMx6* pRS316-*ATG11pr- yeGFP -ATG11*-hphMX6 pRS315-*PHO81pr-pho81(173-1178)-mCherry-his3mx6 (pho81ΔSPX) ade2::ADE2* | This manuscript |
| AG 866 | W303 *Δpho81::kanMx6* pRS316-*prATG11- yeGFP -ATG11-hphMX6* pRS315-*prPHO81-pho81-Y24A-K28A-K154A-mCherry-his3mx6 ade2::ADE2* | This manuscript |
| AG 868 | W303 *Δpho81::kanMx6* pRS316-*prATG11-yeGFP-ATG11-hphMX6* pRS315-*prPHO81-pho81-D453K-D456K-mCherry-his3mx6 ade2::ADE2* | This manuscript |
| ER 75 | PJ69-4a pGAD-C1-*PHO81* pGBDU-C1 | This manuscript |
| ER 77 | PJ69-4a pGAD-C1-*PHO81* pGBDU_C1-*ATG8* | This manuscript |
| ER 79 | PJ69-4a pGAD-C1-*PHO81* pGBDU_C1-*ATG11* | This manuscript |
| AG 73 | W303 *Δatg1::NatMx6* pRS306-*prATG8-2xyeGFP-ATG8 ade2::ADE2* | This manuscript |
| AG 57 | W303 *Δatg11::NatMx6* pRS306-*prATG8-2xyeGFP-ATG8 ade2::ADE2* | This manuscript |
| AG 56 | W303 pRS306-*prATG8-2xyeGFP-ATG8 ade2::ADE2* | This manuscript |
| AG 86 | W303 *Δpho81::TRP1* pRS306-*prATG8-2xyeGFP-ATG8 ade2::ADE2* | This manuscript |
| AG 596 | W303 *ura3:: prADH1-OM45-yeGFP-hphMx6 ade2::ADE2* | This manuscript |
| AG 714 | W303 *Δpho81::Trp1 ura3:: prADH1-OM45-yEGFP- hphMx6 ade2::ADE2* | This manuscript |
| AG 567 | W303 *Pex11-yEGFP-CaURA3 ade2::ADE2* | This manuscript |
| AG 570 | W303 *Δpho81::TRP1 Pex11-yEGFP-CaURA3 ade2::ADE2* | This manuscript |
| AG 716 | W303 *Δatg36:kanMx6 Pex11-yEGFP-CaURA3 ade2::ADE2* | This manuscript |
| AG 718 | W303 *Δatg36:kanMx6 Δpho81::TRP1 Pex11-yEGFP-CaURA3 ade2::ADE2* | This manuscript |
| AG 568 | W303 *Δatg11::natMx6 Pex11-yEGFP-CaURA3 ade2::ADE2* | This manuscript |
| AG 572 | W303 *Δatg11::natMx6 Δpho81::TRP1 Pex11-yEGFP-CaURA3 ade2::ADE2* | This manuscript |
| AG 578 | W303 *Δatg13::natMx6 Pex11-yEGFP-CaURA3 ade2::ADE2* | This manuscript |
| AG 580 | W303 *Δatg13::natMx6 Δpho81::TRP1 Pex11-yEGFP-CaURA3 ade2::ADE2* | This manuscript |

| | | |
|---|---|---|
| AG 937 | W303 *Δnpr2::kanMx6 Pex11-yEGFP-CaURA3 ade2::ADE2* | This manuscript |
| AG 938 | W303 *Δpho81::trp1 Δnpr2::kanMx6 Pex11-yEGFP-CaURA3 ade2::ADE2* | This manuscript |
| AG 75 | W303 *Δatg13::natMx6* pRS306-*prATG8-2xyEGFP-ATG8 ade2::ADE2* | This manuscript |
| AG 80 | W303 *Δatg13::natMx6 Δpho81::kanMx6* pRS306-*prATG8-2xyEGFP-ATG8 ade2::ADE2* | This manuscript |
| AG 917 | W303 *Δatg13::natMx6 ura3::prADH1-OM45-GFP-hphMx6 ade2::ADE2* | This manuscript |
| AG 919 | W303 *Δatg13::natMx6 Δpho81::TRP1 ura3::prADH1-OM45-GFP-hphMx6 ade2::ADE2* | This manuscript |
| ER 232 | PJ69-4a pGAD-C1 pGBDU-C1 | This manuscript |
| ER 234 | PJ69-4a pGAD-C1 pGBDU-C1-*ATG11* | This manuscript |
| ER 236 | PJ69-4a pGAD-C1-*PHO81* pGBDU-C1 | This manuscript |
| ER 238 | PJ69-4a pGAD-C1-*PHO81* pGBDU-C1-*ATG11* | This manuscript |
| ER197 | PJ69-4a pGAD-C1-*pho81-A (PHO81 (410-653)::AKR1 (78-276))* pGBDU-C1-*ATG11* | This manuscript |
| ER199 | PJ69-4a pGAD-C1- *pho81-LA1 (PHO81(455-457)::AKR1(103-107)* pGBDU-C1-*ATG11* | This manuscript |
| ER201 | PJ69-4a pGAD-C1- *pho81-LA2 (PHO81(489-505)::AKR1(138-141))* pGBDU-C1-*ATG11* | This manuscript |
| ER203 | PJ69-4a pGAD-C1- *pho81-LA3 (PHO81(536-555)::AKR1(172-174))* pGBDU-C1-*ATG11* | This manuscript |
| ER205 | PJ69-4a pGAD-C1- *pho81-LA4 (PHO81(587-590)::AKR1(205-212))* pGBDU-C1-*ATG11* | This manuscript |
| ER 240 | PJ69-4a pGAD-C1-*pho81-T461A* pGBDU-C1-*ATG11* | This manuscript |
| ER 242 | PJ69-4a pGAD-C1-*pho81-D453K-D456K* pGBDU-C1-*ATG11* | This manuscript |
| ER 244 | PJ69-4a pGAD-C1-*pho81-D453K-D456K-T461A* pGBDU-C1-*ATG11* | This manuscript |
| ER 246 | PJ69-4a pGAD-C1-*pho81-D806K-D808K* pGBDU-C1-*ATG11* | This manuscript |
| AG 720 | W303 *pRS315 Pex11-yeGFP-CaURA3 ade2::ADE2* | This manuscript |
| AG 722 | W303 *pRS315 Δpho81::TRP1 Pex11-yeGFP-CaURA3 ade2::ADE2* | This manuscript |
| AG 651 | W303 pRS315- *prPHO81-PHO81-mCherry- his3Mx6 Δpho81::TRP1 Pex11-yeGFP-CaURA3 ade2::ADE2* | This manuscript |
| AG 659 | W303 pRS315- *prPHO81-pho81-D453K-D456K-mCherry- his3Mx6 Δpho81::TRP1 Pex11-yeGFP-CaURA3 ade2::ADE2* | This manuscript |
| AG 661 | W303 pRS315- *prPHO81-pho81-T461A-mCherry- his3Mx6 Δpho81::TRP1 Pex11-yeGFP-CaURA3 ade2::ADE2* | This manuscript |
| AG 662 | W303 pRS315- *prPHO81-pho81-D453K-D456K-T461A-mCherry- his3Mx6 Δpho81::TRP1 Pex11-yeGFP-CaURA3 ade2::ADE* | This manuscript |
| AG 848 | W303 pRS315- *prPHO81-pho81(173-1178)-mCherry-his3mx6 (pho81ΔSPX) Δpho81::TRP1 Pex11-yeGFP-CaURA3 ade2::ADE2* | This manuscript |
| AG 849 | W303 pRS315- *prPHO81-pho81-Y24A-K28A-K154A-mCherry- his3Mx6 Δpho81::TRP1 Pex11-yeGFP-CaURA3 ade2::ADE2* | This manuscript |
| AG 851 | W303 *pRS315 Δatg13::natMx6 Δpho81::TRP1 Pex11-yeGFP-CaURA ade2::ADE2* | This manuscript |
| AG 852 | W303 pRS315-*prPHO81-PHO81-mCherry- his3Mx6 Δatg13::natMX6 Δpho81::TRP1 Pex11-yeGFP-CaURA3 ade2::ADE2* | This manuscript |
| AG 855 | W303 pRS315- *prPHO81-pho81-D453K-D456K-T461A-mCherry- his3Mx6 Δatg13::natMx6 Δpho81::TRP1 Pex11-yeGFP-CaURA3 ade2::ADE2* | This manuscript |
| AG 856 | W303 pRS315-*prPHO81-PHO81-mCherry- his3Mx6* pRS316 *Δatg8::GFP-natMx6* | This manuscript |
| AG 862 | W303 pRS316-*prATG11-yeGFP-ATG11-hpMx6* pRS315-*prPHO81-PHO81-mCherry- his3Mx6 Δpho81::kanMx6 ade2::ADE2* | This manuscript |
| AG 868 | W303 pRS316-*prATG11-yeGFP-ATG11-hpMx6* pRS315-*prPHO81-pho81-D453K-D456K-mCherry- his3Mx6 Δpho81::kanMx6 ade2::ADE2* | This manuscript |

| | | |
|---|---|---|
| AG 864 | W303 pRS316-*prATG11-yeGFP-ATG11-hpMx6* pRS315- *prPHO81-pho81(173-1178)-mCherry-his3mx6 (pho81ΔSPX) Δpho81::TRP1 ade2::ADE2* | This manuscript |
| AG 866 | W303 pRS316-*prATG11-yeGFP-ATG11-hpMx6* pRS315- *prPHO81-pho81-Y24A-K28A-K154A-mCherry-his3mx6 Δpho81::TRP1 ade2::ADE2* | This manuscript |
| YMG1319, RG619 | W303 pRS315-*prADH1-PHO81-yEGFP-caURA3 Δpho81::natMx6* | This manuscript |
| YMG1322, RG622 | W303 pRS315-*prADH1-pho81(173-1178)-yEGFP-caURA3 (pho81ΔSPX) Δpho81::natMx6* | This manuscript |
| YMG1323, RG623 | W303 pRS315-*prADH1-pho81(1-172)-yEGFP-caURA3 (pho81-SPXonly) Δpho81::natMx6* | This manuscript |
| AG 881 | W303 pRS315- *prPHO81-pho81(173-1178)-mCherry- his3Mx6 (pho81ΔSPX) Δatg13::natMx6 Δpho81::TRP1 Pex11-yeGFP-CaURA3 ade2::ADE2* | This manuscript |
| AG 883 | W303 pRS315- *prPHO81-pho81-Y24A-K28A-K154A-mCherry- his3Mx6 Δatg13::natMx6 Δpho81::TRP1 Pex11-yeGFP-CaURA3 ade2::ADE2* | This manuscript |
| AG 874 | W303 pRS315 *Δpho4::kanMx6 Pex11-yeGFP-CaURA3 ade2::ADE2* | This manuscript |
| AG 875 | W303 pRS315 *Δpho4::kanMx6 Δpho81::TRP1 Pex11-yeGFP-CaURA3 ade2::ADE2* | This manuscript |
| AG 876 | W303 pRS315-*prPHO81-PHO81-mCherry- his3Mx6 Δpho4::kanMx6 Δpho81::TRP1 Pex11-yeGFP-CaURA3 ade2::ADE2* | This manuscript |
| AG 877 | W303 pRS315-*prPHO81-pho81-D453K-D456K-mCherry- his3Mx6 Δpho4::kanMx6 Δpho81::TRP1 Pex11-yeGFP-CaURA3 ade2::ADE2* | This manuscript |
| AG 878 | W303 pRS315-*prPHO81-pho81(173-1178)-mCherry- his3Mx6* (*pho81ΔSPX*) *Δpho4::kanMx6 Δpho81::TRP1 Pex11-yeGFP-CaURA3 ade2::ADE2* | This manuscript |
| AG 879 | W303 pRS315-*prPHO81-pho81-Y24A-K28A-K154A-mCherry-his3Mx6 Δpho4::kanMx6 Δpho81::TRP1 Pex11-yeGFP-CaURA3 ade2::ADE2* | This manuscript |
| AG 160 | pRS315-*prADH1-PHO81-mCherry-His3mx6 Δpho81::TRP1* pRS306-*prATG8-2xyeGFP-ATG8 ade2::ADE2* | This manuscript |
| AG 762 | W303 *Δpho91::TRP1 Pex11-yeGFP-CaURA3 ade2::ADE2* | This manuscript |
| AG 870 | W303 *Δpho4::kanMx6 Pex11-yeGFP-CaURA3 ade2::ADE2* | This manuscript |
| AG 871 | W303 *Δpho4::kanMx6 Δpho81::TRP1 Pex11-yeGFP-CaURA3 ade2::ADE2* | This manuscript |
| AG 724 | W303 pRS315-*prADH1-PHO81-mCherry-His3mx6 Δpho81::TRP1 Pex11-yeGFP-CaURA3 ade2::ADE2* | This manuscript |
| AG 872 | W303 pRS315-*prCUP1-PHO81-mCherry-His3mx6 Δpho81::TRP1 Pex11-yeGFP-CaURA3 ade2::ADE2* | This manuscript |
| AG 873 | W303 pRS315-*prPHO5-PHO81-mCherry-His3mx6 Δpho81::TRP1 Pex11-yeGFP-CaURA3 ade2::ADE2* | This manuscript |
| AG 161 | W303 pRS315-*prADH1-pho81-A(PHO81 (410-653)::AKR1 (78-276))-mCherry-His3mx6 Δpho81::TRP1* pRS306-*prATG8-2xyeGFP-ATG8 ade2::ADE2* | This manuscript |
| AG 149 | W303 pRS315-*prADH1-pho81-LA1(PHO81(455-457)::AKR1(103-107) -mCherry-His3mx6 Δpho81::TRP1* pRS306-*prATG8-2xyeGFP-ATG8 ade2::ADE2* | This manuscript |
| AG 150 | W303 pRS315-*prADH1-pho81-LA2(PHO81(489-505)::AKR1(138-141)) -mCherry-His3mx6 Δpho81::TRP1* pRS306-*prATG8-2xyeGFP-ATG8 ade2::ADE2* | This manuscript |
| AG 277 | W303 pRS315-*prADH1-pho81-LA3(PHO81(536-555)::AKR1(172-174))-mCherry-His3mx6 Δpho81::TRP1* pRS306-*prATG8-2xyeGFP-ATG8 ade2::ADE2* | This manuscript |
| AG 152 | W303 pRS315-*prADH1-pho81-LA4 LA4 (PHO81(587-590)::AKR1(205-212))-mCherry-His3mx6 Δpho81::TRP1* pRS306-*prATG8-2xyeGFP-ATG8 ade2::ADE2* | This manuscript |
| AG 310 | W303 *Δatg19::HIS3 Δtrp1::mcherry-ATG8 ATG1-3xyeGFP-caUra* | This manuscript |
