## Supplementary material for "A metabolite sensor subunit of the Atg1/ULK complex regulates selective autophagy": Table S2

Table S2: Differential expression analysis and statistics performed using limma

| Fig.4f | GFP-Atg11/CTRL with Pho81 expression | | | GFP-Atg11/CTRL with Pho81$^{DD}$ expression | | |
|---|---|---|---|---|---|---|
| Gene name | adj. P Value | $log_2$ FC | significant | adj. P Value | $log_2$ FC | significant |
| PHO81 | 9.39E-06 | 5.949 | + | 2.96E-03 | -2.92 | + |
| ATG1 | 3.10E-05 | 9.67 | + | 1.12E-04 | 10.178 | + |
| ATG9 | 4.27E-05 | 6.121 | + | 5.27E-04 | 5.303 | + |
| ATG11 | 3.57E-06 | 16.787 | + | 1.05E-05 | 17.041 | + |
| ATG13 | 2.24E-05 | 8.413 | + | 1.74E-04 | 7.92 | + |
| ATG17 | 8.37E-06 | 12.446 | + | 3.35E-05 | 12.572 | + |
| ATG19 | 1.35E-05 | 11.34 | + | 4.50E-05 | 12.261 | + |
| ATG29 | 5.56E-06 | 11.788 | + | 2.34E-05 | 11.731 | + |
| ATG31 | 9.39E-06 | 11.782 | + | 4.50E-05 | 11.827 | + |
| ATG32 | 3.68E-03 | 1.791 | + | 3.53E-01 | -0.536 | |
| ATG34 | 8.98E-06 | 9.874 | + | 4.22E-04 | 6.563 | + |
| ATG36 | 2.45E-04 | 3.337 | + | 4.06E-03 | 2.682 | + |

| Fig. 5c | GFP-Atg11/CTRL with Pho81 expression | | | GFP-Atg11/CTRL with Pho81$^{\Delta SPX}$ expression | | | GFP-Atg11/CTRL with Pho81$^{YKK}$ expression | | |
|---|---|---|---|---|---|---|---|---|---|
| Gene name | adj. P Value | log2FC | significant | adj. P Value | log2FC | significant | adj. P Value | log2FC | significant |
| PHO81 | 1.35E-07 | 4.786 | + | 2.97E-08 | 5.568 | + | 2.38E-08 | 5.671 | + |
| ATG1 | 5.85E-11 | 9.157 | + | 5.90E-11 | 9.15 | + | 8.32E-11 | 8.902 | + |
| ATG11 | 1.81E-15 | 16.112 | + | 1.95E-15 | 16.023 | + | 2.61E-15 | 15.653 | + |
| ATG13 | 1.40E-01 | 1.822 | | 2.24E-02 | 3.132 | + | 7.64E-02 | 2.047 | |
| ATG17 | 2.79E-08 | 8.35 | + | 4.16E-09 | 10.044 | + | 2.53E-08 | 8.472 | + |
| ATG19 | 1.02E-12 | 9.932 | + | 3.80E-12 | 8.942 | + | 1.80E-12 | 9.493 | + |
| ATG29 | 2.70E-09 | 10.248 | + | 3.08E-09 | 10.163 | + | 3.20E-09 | 10.131 | + |
| ATG31 | 6.31E-12 | 10.225 | + | 5.56E-12 | 10.329 | + | 1.23E-11 | 9.694 | + |
| ATG34 | 3.15E-01 | 1.258 | | 2.61E-03 | 4.62 | + | 2.37E-01 | 1.342 | |
