## Supplementary figures and images for "A metabolite sensor subunit of the Atg1/ULK complex regulates selective autophagy"

### Extended data 1

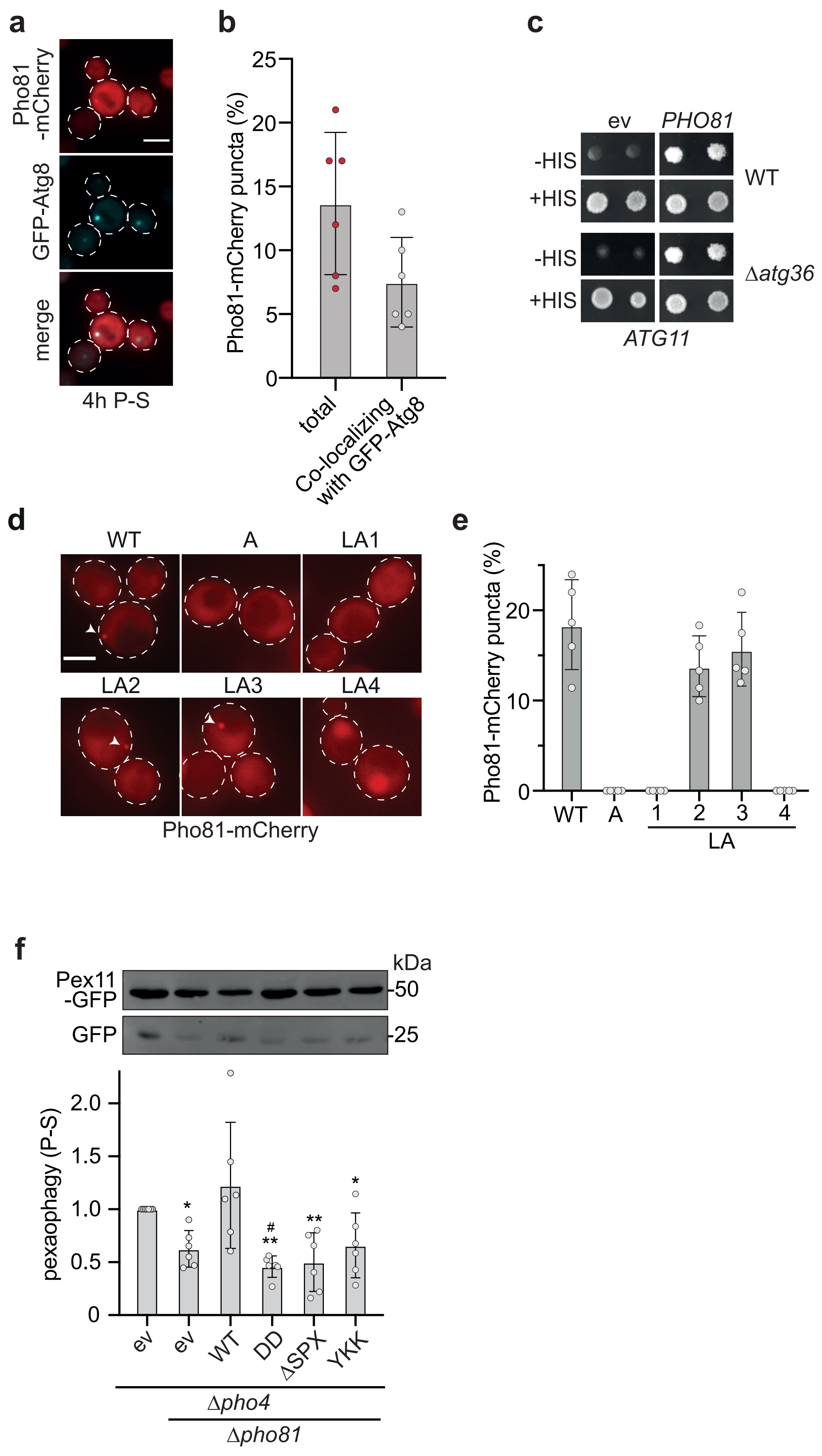


**Extended Data 1**
